# Unbiased data-driven analysis of five amyloid-beta peptides for biomarker investigations in familial Alzheimer’s disease

**DOI:** 10.1101/2024.11.23.624811

**Authors:** Isaac Llorente-Saguer, Rebecca Gabriele, Teisha Bradshaw, Claire A. Leckey, Christopher R.S. Belder, Lucía Chávez-Gutiérrez, Rohan de Silva, Nick C. Fox, Selina Wray, Neil P. Oxtoby, Charles Arber

## Abstract

**INTRODUCTION:** Changes to the relative abundance of amyloid-beta (Aβ) peptides are hallmarks of Alzheimer’s disease (AD). iPSC-derived neurons offer a physiological model of Aβ production. We employed unbiased, data-driven analyses to investigate combinations of Aβ peptides as AD biomarkers and the relative contribution of peptides to AD pathogenesis.

**METHODS:** We measured Aβ37, Aβ38, Aβ40, Aβ42 and Aβ43 in ten iPSC-neuronal cultures from *PSEN1* mutation carriers. We combined these data with published cell model data and used linear weighted combinations to 1) distinguish AD from controls, and 2) predict age-at-onset for *PSEN1* mutations.

**RESULTS:** Data-driven approaches distinguished Aβ42 and Aβ43 from shorter peptides, providing unbiased evidence for their contribution to disease. Weighted linear combinations of Aβ peptides outperform Aβ42/40 and provide insights into relative peptide contribution as biomarkers; the optimal ratio for all data is represented as (21 · Aβ37 + 10 · Aβ38 + 69 · Aβ40)/(94 · Aβ42 + 6 · Aβ43).

**DISCUSSION:** The algorithm discovered herein can be further refined to improve biomarkers for AD.

## 1. Background

Mutations in *PSEN1* that cause early onset familial Alzheimer’s disease (EOFAD) reduce the processing of amyloid-beta (Aβ) from longer peptides (e.g., Aβ42 and Aβ43) to shorter peptides (e.g., Aβ37, Aβ38 and Aβ40)^1–3^. This causes a relative increase in longer, more aggregation-prone Aβ species, potentially predisposing Aβ aggregation, plaque pathology and disease^4^. For this reason, a raised Aβ42/40 ratio is used in model systems to validate pathogenicity of EOFAD causing mutations^5^, and is also supported by clinical data in plasma of mutation carriers^6^ (although, a recent study did not replicate these changes in plasma^7^). This phenomenon is distinct from the reduced Aβ42:40 ratio observed in plasma and CSF that is used to establish the presence of Aβ pathology in both EOFAD and sporadic AD (possibly due to higher sequestration of Aβ42 within plaques in the brain parenchyma).

Recently, Liu and colleagues reported that the Aβ37/42 ratio represents an improved biomarker for AD compared to Aβ42/40^8^. In parallel work, a ratio of Aβ37+38+40/Aβ42+43 (termed “Aβshort/long” hereafter) was shown to accurately predict the age-at-onset of EOFAD mutations in cell models^9,10^. Together, these studies show the value of incorporating a diversity of Aβ peptides into biomarker studies and prompt detailed exploration of peptide combinations as biomarkers.

*PSEN1* mutation carriers show remarkable heterogeneity in terms of age at onset^11^, clinical symptoms^12^ and ratios of Aβ peptides^10,13^. Indeed, certain *PSEN1* mutations significantly affect the Aβ43/40 ratio whilst leaving the Aβ42/40 ratio relatively spared^13–16^. Differences in the relative abundance of Aβ peptides and clinical heterogeneity make it hard to interpret the relative importance of the different Aβ species, their contribution to disease mechanisms and their value as biomarkers.

Induced pluripotent stem cell (iPSC)-derived neurons offer a physiological human neuronal model of Aβ production that faithfully represent Aβ production *in vitro*^13^. Importantly, iPSC-neurons also express one healthy *PSEN1* allele, in contrast to the cell models previously employed^8–10^. Given the similarity of iPSC-neuronal Aβ profiles with blood plasma ratios from the same donor^6^, coupled with the current interest in blood-based biomarker testing for AD, we sought to investigate Aβ ratios in our physiological model and employ unbiased data-driven techniques to probe their use for 1) distinguishing EOFAD from control samples and 2) predicting age-at-onset for EOFAD mutations.

## 2. Methods

### 2.1 Cell culture

Details of all iPSC lines are presented in Table 1. *PSEN1* E184D iPSCs were reprogrammed from fibroblasts for this study (Fig S1) using episomal reprogramming^17^. For the newly established E184D iPSC line, Sanger sequencing was performed (Source Bio) to demonstrate the presence of the mutation (Fig S1B) and a stable karyotype was confirmed using the Stem Cell Technologies hPSC Genetic Analysis kit (Fig S1C). Low coverage whole genome sequencing further confirmed a stable karyotype of the newly generated E184D patient-derived line (Fig S1D)^18^. iPSCs were grown in Essential 8 media on Geltrex substrate and passaged using EDTA mechanical passaging. iPSCs were differentiated into cortical neurons following published protocols^13–15,19^ using dual SMAD inhibition and extended neurogenesis. The final timepoint was taken as 100 DIV, where iPSC-neuronal media was conditioned for 48 hours prior to Aβ analysis.

**Table 1.** iPSC lines used in this study.

| Line | Details and reference | Female/Male | Age at biopsy | APOE status | Age at onset |
| --- | --- | --- | --- | --- | --- |
| Control 1 | KOLF2.1J<sup>20</sup> | Male | 55-59 | 3/3 | - |
| Control 2 | ND41886<sup>13</sup> | Male | 64 | 2/3 | - |
| Control 3 | RBi01 (Sigma)<sup>13</sup> | Male | 45-49 | 3/3 | - |
| *PSEN1* int4del | EBiSC<sup>13</sup> | Female | 47 | 3/3 | 47 |
| *PSEN1* Y115H | *PSEN1* Y115H<sup>13</sup> | Male | 39 | 3/3 | 34 |
| *PSEN1* M139V | EBiSC<sup>13</sup> | Female | 45 | 2/3 | 34 |
| *PSEN1* M146I | EBiSC<sup>13</sup> | Male | 38 | 3/3 | - |
| *PSEN1* E184D | This study | Female | 43 | 3/3 | 45 |
| *PSEN1* R278I | *PSEN1* R278I cA<sup>13</sup> | Male | 60 | 2/4 | 58 |
| *PSEN1* E280G #1 | *PSEN1* E280G.A<sup>14</sup> | Male | 45 | 3/3 | 41 |
| *PSEN1* E280G #2 | *PSEN1* E280G.B<sup>14</sup> | Male | 49 | 3/4 | 38 |
| *PSEN1* P436S #1 | *PSEN1* P436S #1<sup>15</sup> | Female | 52 | 2/3 | 48 |
| *PSEN1* P436S #2 | *PSEN1* P436S #2<sup>15</sup> | Female | 48 | 3/3 | 44 |

### 2.2 Aβ ELISAs

Aβ42, Aβ40 and Aβ38 were quantified using electrochemiluminescence (Meso Scale Discovery, 6E10 V-Plex). Aβ43 was quantified using the IBL Amyloid Beta (1-43) ELISA.

To develop a custom Aβ37 ELISA, we used the β-amyloid 1-37 (D2A6H) antibody from Cell Signaling Technology (carrier-free). Capture antibody was adsorbed to plates at 1mg/ml (empirically optimised) using 0.05M carbonate buffer (Sigma) for 24 hours. Plates were washed and blocked with Starting Block (Thermo) before adding 50μl of undiluted conditioned media or Aβ37 standards (ANASPEC, 0-500pg/ml) in 20% SuperBlock (Thermo) together with 50μl of capture antibody 6E10-HRP (Biolegend, 3mg/ml (empirically optimised)). After washing, colour development was initiated with TMB blue reagent and stopped using 2M sulphuric acid. Absorbance was measured at 450nm on a Tecan Spark 10M plate reader.

Other cell data used in this study include published Aβ profiles generated by overexpression of fAD-linked *PSEN1* mutations in *PSEN1*/*PSEN2* double knockout cell lines; mouse embryonic fibroblast cell lines by Petit and colleagues^9^, and by overexpression in HEK 293T cell lines by Liu and colleagues^8^ and Schultz and colleagues^10^.

For iPSC data, actual age-at-onset for each donor was used where available, otherwise, age-at-onset was predicted from individuals from within families with the same mutation as the donor^12^. For Liu, Schultz and Petit data age-at-onset was taken from data from within each publication.

Modelling was performed on the mean ratio for each individual patient-derived line or genotype.

### 2.3 Modelling/Analysis

#### Data harmonisation

Samples with any peptide measure equal to zero were removed (N = 14 from a total of 230; 3 from Petit et al.^9^, and 11 from Liu et al.^8^). Zeros could potentially be due to assay detection limits or errors, and impede the usage of the peptide as a denominator, for these samples. Samples were averaged across batches to reduce the influence of measurement error. Data harmonisation across all datasets was performed in two steps. First, each sample’s data were divided by the sum of the available peptides within that sample. This step adjusts for differences in total peptide amounts across samples, as we are interested in their relative amounts. Second, each peptide measure was further divided by the average of the control samples for the same dataset to correct for any assay-specific systematic biases.

To evaluate biomarkers that include the addition of multiple peptides, each peptide measurement was scaled back to the relative control proportions based on data from Petit et al^9^. Ratio biomarkers like the short/long^9^ involve a weighted sum where each peptide is weighted by its naturally occurring abundance in controls. This scaling step is essential because the relative abundances of Aβ peptides differ by orders of magnitude. Not scaling the data back will distort the setup for which the biomarker was designed.

#### Statistical analysis

We evaluate biomarkers in case-control classification and age-at-onset regression scenarios. In classification experiments we analyse the area under the receiver operator characteristic curve (ROC AUC), which plots sensitivity, or true positive rate TP/(TP+FN) against false positive rate FP/(FP+TN), where T/F indicates True/False and P/N indicates Positive/Negative. Since the data are strongly unbalanced (very few control samples), we also look at the precision-recall curve (PR AUC) of the smaller group, which plots control precision TN/(TN+FN) against control recall TN/(TN+FP), since PR AUC is a preferred metric in this scenario.

To evaluate the correlation between the biomarker and the age-at-onset (AAO), we employ the coefficient of determination (R^2^), using Pearson correlation, since peptide biomarkers have shown good linear correlation^9^.

Median and 95% confidence intervals are reported for all metrics using 2000 bootstrapping samples, stratified in the classification case (to accommodate the heavy class imbalance).

Finally, we bootstrap the data-driven combinatorial biomarker “weighted composite value ratio” (wCVR, explained below), running the search algorithm three times per bootstrap with random initialisations, to investigate the effect of the weighting of the peptides. We also do a grid-search analysis, where we fixed the Aβ42/40 ratio as baseline for R^2^, and then calculated the metrics of all combinations of the other peptide weights from 0 to 1 by 0.02 (either in the numerator or denominator). We did the same with precision-recall of the controls, but the baseline ratio was Aβ37/42.

#### Biomarker combinations

Different biomarker combinations have been proposed in the field, namely Aβ42/40, Ab37/42 and the short/long ratio, which is Aβ(37+38+40)/(42+43). We ran BioDisCVR^21^ to find which peptides are selected to go to the numerator, and which are selected to go to the denominator, along with their linear combination weights, obtaining a weighted combined value ratio (wCVR). The general formula is

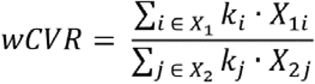

, where X_1_ and X _2_ are non-overlapping non-empty sets of the available peptides, and k is a weighting factor for each peptide. To drive the algorithm, we consider different configurations, which we indicate with the suffixes -R for R^2^, -P for PR AUC, and RP for both (i.e., when the objective function is the product of both metrics). The BioDisCVR framework allows for a user-chosen search algorithm. We chose to run a genetic algorithm^22^, as described in the original paper.

Additionally, we evaluated all possible combinations of ratios (180 possible combinations) where either the numerator or denominator is one peptide, or the addition of multiple peptides, without repetition, and conserving the different proportions of the peptides from Petit et al.

## 3. Results

We established a novel Aβ37 ELISA that showed linearity, specificity and was not sensitive to matrix effect (Fig S2). We observed only minimal Aβ38 cross-reactivity, producing a signal that was 34-fold lower than the Aβ37 specific signal. We expect similar concentrations of Aβ37 and Aβ38^8,13^, meaning the contribution of non-specific signal is negligible. Importantly, our results are comparable with our published mass spectrometry findings from the same iPSC-neuronal lines (Fig S2D).

Figure 1 shows the proportion of Aβ37, Aβ38, Aβ40, Aβ42 and Aβ43 measured in conditioned media from 10 *PSEN1* mutation carrying patient-derived iPSC-neuronal lines in triplicate, as well as 3 controls. As expected for EOFAD data, there is visual separation between cases and controls, as well as variation between *PSEN1* patient-derived lines.

**Figure 1.**
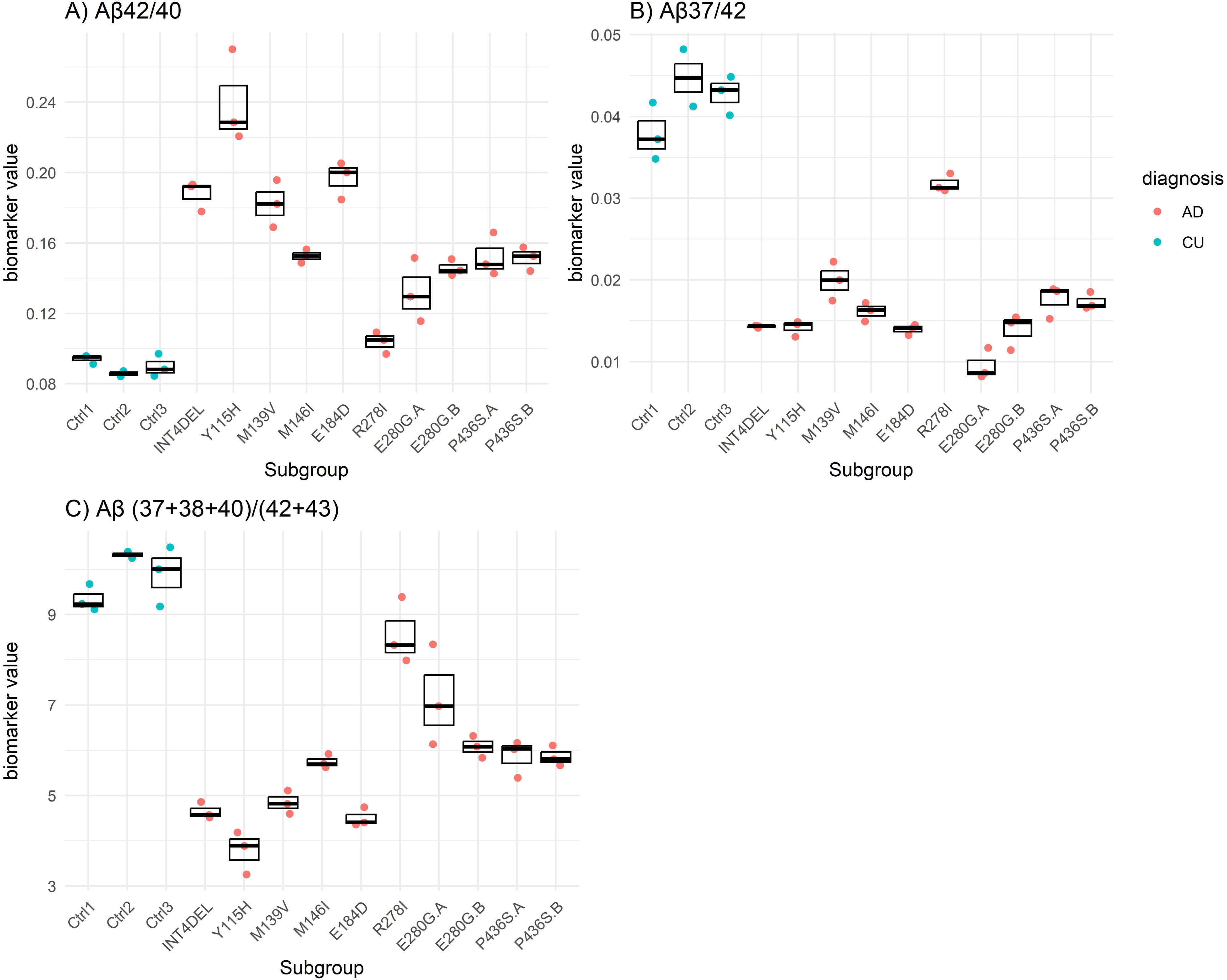
Aβ ratios in iPSC data. Each data point represents an independent neuronal differentiation from each iPSC line (n=3 throughout, apart from Ctrl2 where n=2). Control iPSC-derived neurons and *PSEN1* patient-derived iPSC-neurons are shown in blue and red respectively.

To allow comparison of Aβ data from published datasets and our newly generated data, harmonisation was required because of widely acknowledged inter-assay variability (Figure 2). Harmonisation was achieved by weighting peptides based on the average relative abundance of each peptide found in control cell data within each dataset. Converting to a dimensionless, normative scale (Figure 2 right panels) facilitates both data interpretation and combination of multi-assay datasets (Aβ peptide ratios have been shown to be highly replicable between control lines^13^). Figure S3 shows the relative peptide amount of control cell lines for each dataset before harmonisation. Although all datasets share similar general relative abundance (e.g., Aβ40 being the most abundant, and Aβ43 being the least), there are some important differences, especially for our cohort (iPSC), where the relative abundance of Aβ37 is lower and the Aβ38 is higher than the rest of the cohorts.

**Figure 2.**
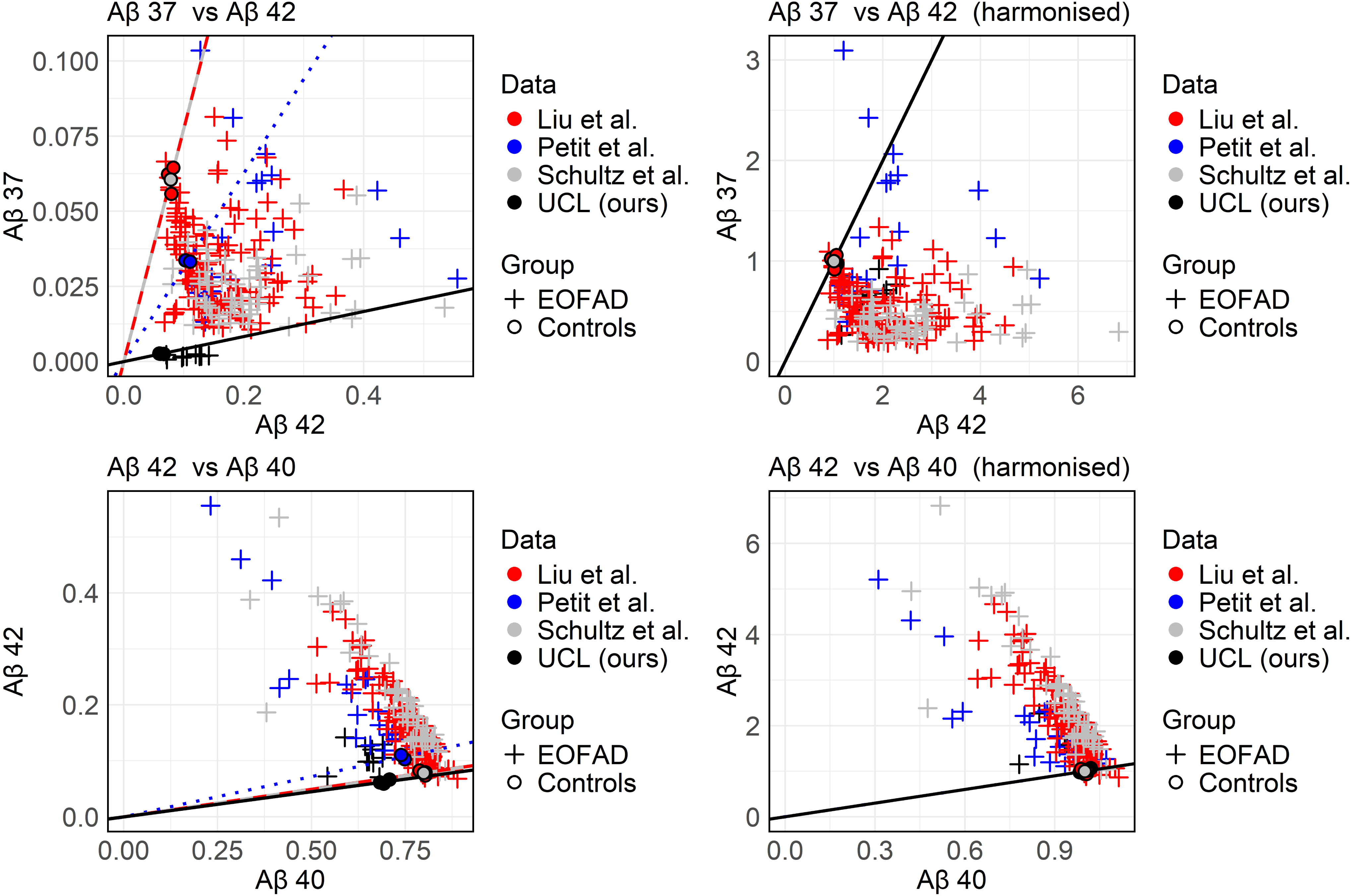
Data harmonisation and peptide distribution. A) Left: raw data, divided by total peptide count. Right: data harmonised by dividing by assay-specific average in controls. Top row: Aβ37 vs Aβ42. Lower row: Aβ42 vs Aβ40. Datasets are colour coded. Lines depict the direction of data from controls, showing the successful harmonisation.

Figure 3 shows the peptide distribution of all EOFAD data, relative to the controls. Most deviations of Aβ37, Aβ38 and Aβ40 occur below the mean of controls (91.3%, 82.6% and 72.9%, respectively), while most deviations of Aβ42 and Aβ43 occurred above the mean of controls (97.1% and 84.5%, respectively).

**Figure 3.**
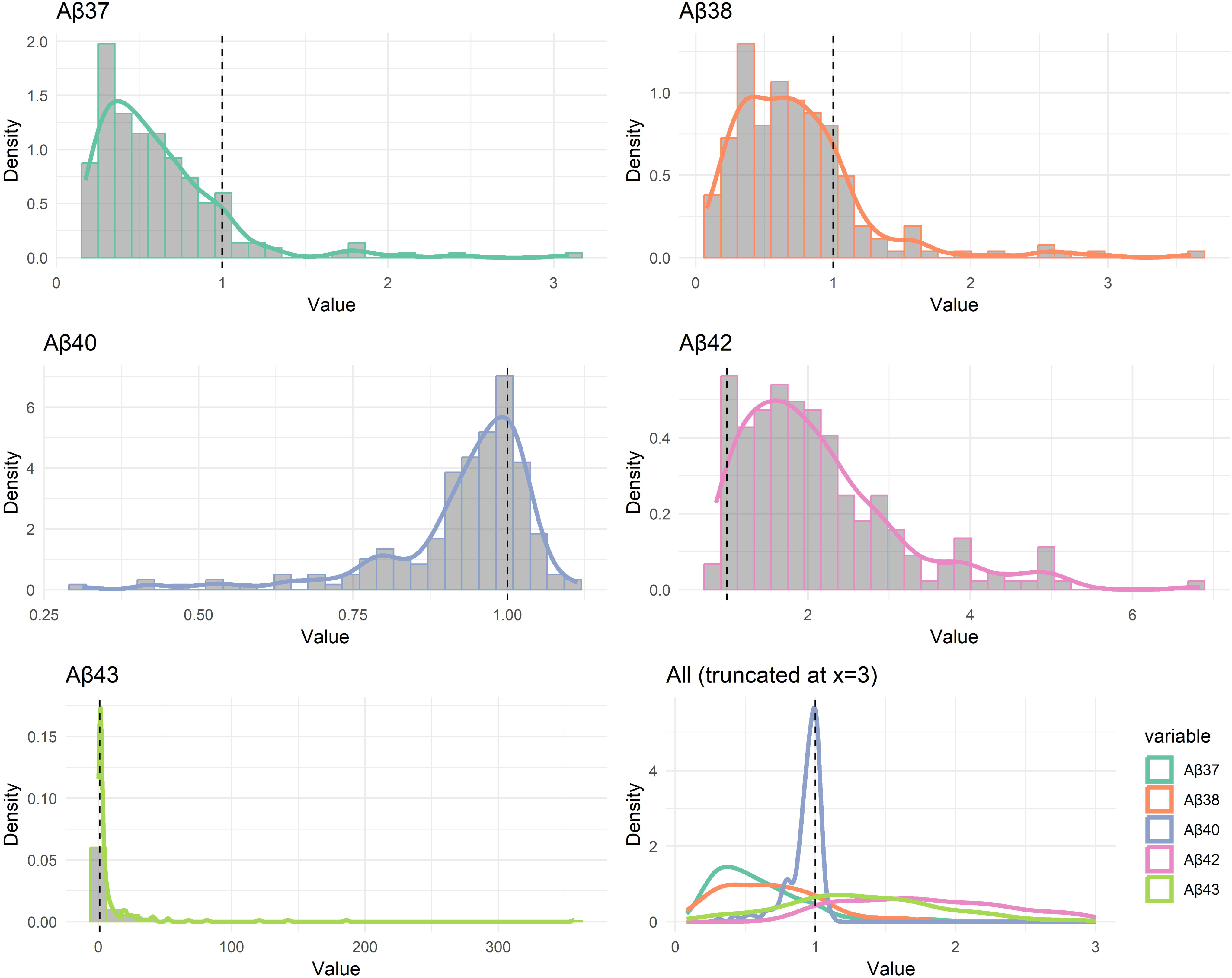
Aβ peptide distribution from EOFAD data relative to controls. Representation of all EOFAD data deviating from the mean of controls, set as 1. Each Aβ peptide distribution is shown, plus all peptides combined (lower right). Most deviations of Aβ37, Aβ38 and Aβ40 occur below the controls (91.3%, 82.6% and 72.9%, respectively), while deviations of Aβ42 and Aβ43 occurred above the mean of controls (97.1% and 84.5%, respectively).

Table 2 shows the correlation (Pearson and Kendall) with the age-at-onset for all peptides, per dataset (pre-transformation), and for all data merged (post-transformation). The measures are all relative to the sum of the five peptides. All the statistically significant comparisons (p < 0.05, in bold, and thus the subset of corrected p-values) share the same message: positive correlation for Aβ37, Aβ38 and Aβ40, and negative correlation for Aβ42 and Aβ43.

Using all harmonised data, we assessed the association between the position of the mutation in *PSEN1* and age-at-onset. The Kendall’s τ coefficient was calculated to be 0.0104 (z = 0.228, p = 0.820). The Pearson correlation coefficient was 0.0118 (t = 0.175, df = 219, p-value = 0.861). The practically zero coefficients and high p-values suggest that the observed correlation is not statistically significant. Figure S4 shows a scatterplot of AAO versus location, and Aβ42/40 versus location.

A dataset-specific evaluation of biomarkers is shown as heatmaps in Figure 4. We compare established ratios (Aβ42/40, Aβ40/42, Aβ37/42 and Aβshort/long) with weighted combined value ratio (wCVR) analyses with different configurations (see methods), where wCVR was optimised per cohort. For both correlation with age-at-onset (coefficient of determination, Figure 4, left panel) and case-control classification (precision-recall, Figure 4, right panel), the heatmaps show performance heterogeneity between the cohorts. Data from iPSC, Petit or Schultz classify well (perfect classification for all biomarkers except for Aβ42/40, Aβ37/42 and Aβ40/42), and data from Petit exhibits highest coefficient of determination, followed by iPSC. As for the novel biomarkers, wCVR achieves superior performance within each metric to which it is directed, namely wCVR-P achieves the best classification for precision recall of the controls, and wCVR-R shows the best correlation with age-at-onset when driven by the coefficient of determination. Aβshort/long and Aβ40/42 also perform well for correlation with age-at-onset.

**Figure 4.**
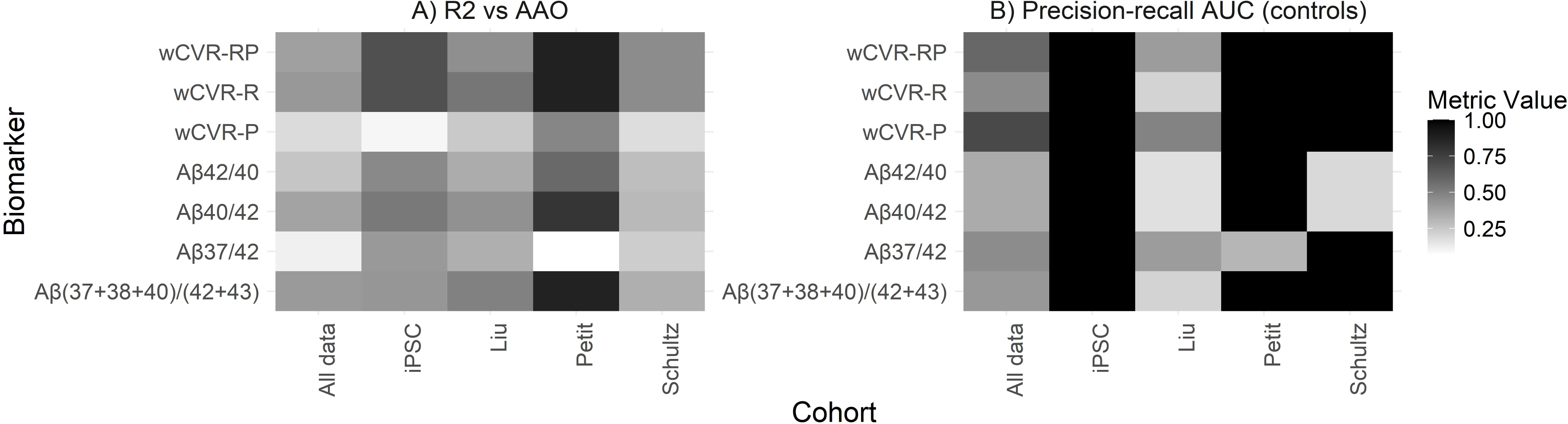
Biomarker performance summary. Left: age-at-onset correlation, measured by the coefficient of determination, R^2^. Right: case-control classification, measured by area under the precision-recall curve. Rows show biomarker performance (increasing grayscale) for each cohort/dataset (columns), with wCVR performing well on both tasks across all datasets. Abbreviations: AAO – age-at-onset. wCVR – weighted composite value ratio, where P = precision recall (disease classification), R = R2 (coefficient of determination), and RP = both.

Figure 5 shows the results of our ratio biomarker experiments. The map of R^2^ (AAO) vs precision-recall AUC shows wCVR biomarkers towards the upper right quadrant, demonstrating superior performance to most simple combinations of peptides. The literature biomarkers also do well in one metric each, but not both: Aβshort/long shows high R^2^, and Aβ37/42 outperforms most simple ratios in precision-recall AUC.

**Figure 5.**
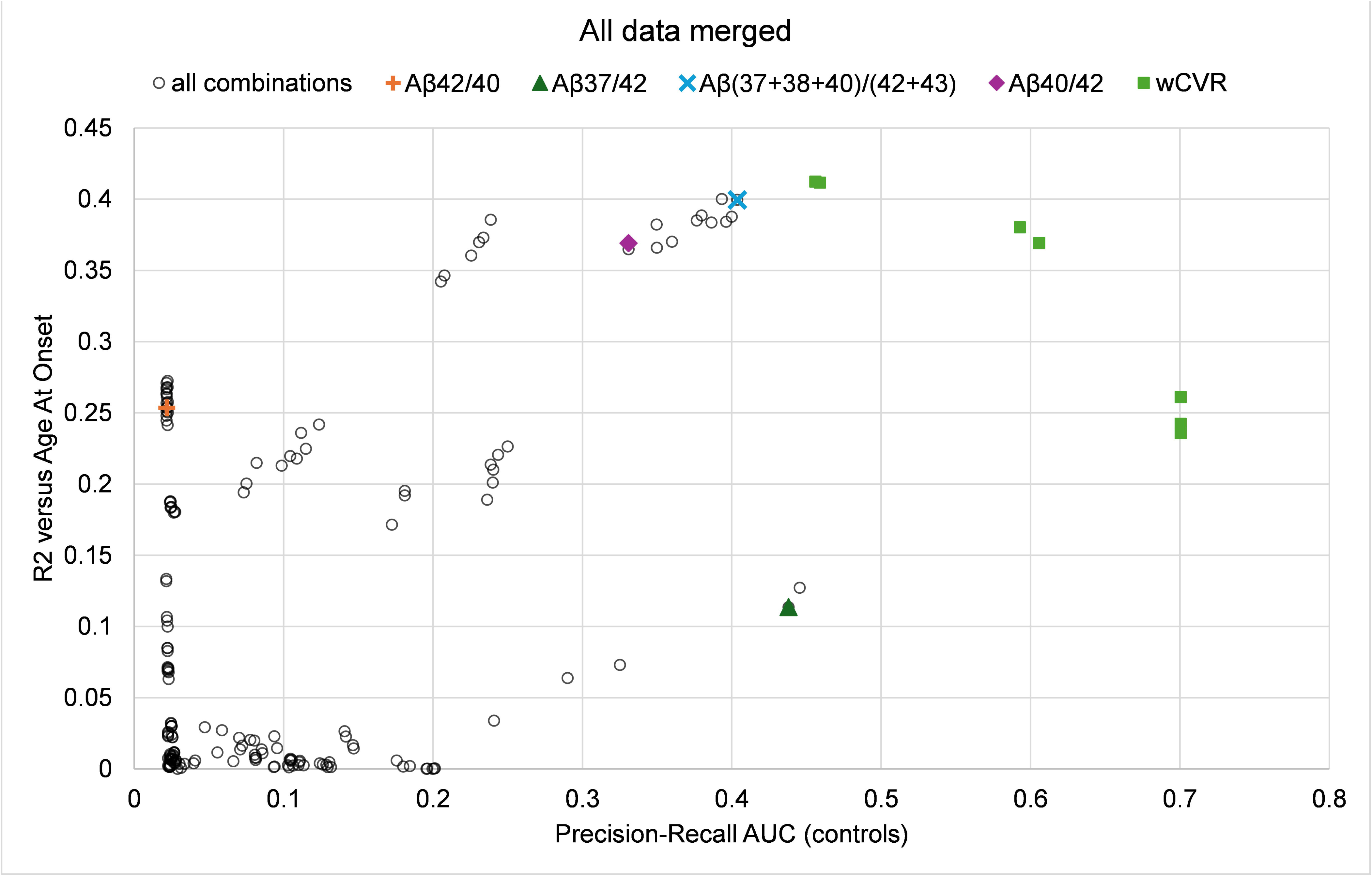
Exploration of ratio biomarker performance. Scatter plot showing performance on both AAO correlation (R^2^, vertical axis) and case-control classification (precision-recall AUC, horizontal axis). Of all simple ratios (empty circles), some show good performance in one metric or the other. wCVR biomarkers consistently show strong performance in both metrics (upper right quadrant).

Table 3 contains the quantification of the performance of the salient biomarkers, with 95% confidence intervals from bootstrapping (see Methods). It is noteworthy that wCVR is able to improve on established ratios from the literature, however, different wCVR biomarkers excel at precision-recall versus coefficient of determination.

Table 4 lists the peptide weights (denominator in red and negative numbers) for the different wCVR configurations, along with their classification and regression performance. There is a trade-off between the two evaluation metrics (classification and regression), which is determined by the weight shifting and position in the ratio of the peptides. In all cases, Aβ40 is in the numerator and Aβ42 and Aβ43 are in the denominator. Aβ38 is actively (10∼19%) in the numerator when the objective function includes the coefficient of determination, and Aβ37 appears in the numerator when classification (precision-recall) is considered. When considering both metrics of classification and regression, the composite value ratio becomes similar to the short/long ratio, with notable more weighting of the Aβ42 in the denominator, and Aβ40 in the numerator. Table 4B shows the wCVR for the individual datasets, demonstrating the different contribution of each peptide to combined performance (precision-recall plus coefficient of determination) for each dataset. Note that Aβ42 and Aβ43 are consistently in the denominator, while Aβ37, Aβ38 and Aβ40 are in the numerator.

Results from a permutation feature importance analysis are shown in Supplementary Table S1. For all metrics, and for both displayed biomarkers (Aβshort/long and wCVR-RP), Aβ42 is the peptide providing the biggest impact. The data-driven wCVR-RP better employs information from Aβ43, as the importance values are higher in all metrics.

Different patterns were observed, depending on the metric used for optimisation, when bootstrapping the aggregated data and running BioDisCVR three times per bootstrap to get the best wCVR. Figure S5 shows the boxplots of the weights for 200 bootstrapped samples, where a positive weight places the peptide in the numerator, and a negative weight in the denominator of the ratio. However, these weights were obtained by fitting different resampled data, so to better understand their effect, Figure S6 shows the evaluation metrics against peptide weights. In all cases, we can observe a clear convex space of data points, with a slight truncation for the precision-recall plots. A similar pattern of an optimal weight combination can be observed in Figure S7, where the grid-search analysis of weighted combinations shows a single peak of performance (yellow). These data suggest that employing multiple peptides improves performance in each case.

## 4. Discussion

We measured five neuronal Aβ peptides from 10 patient-derived iPSC lines to investigate Aβ ratios in an unbiased, data-driven manner. The data independently verify a differential contribution of longer peptides (Aβ42 and Aβ43) and shorter peptides (Aβ40, Aβ38 and Aβ37) to disease. We explored the relative contribution of each peptide to disease classification and onset prediction. Aβ40 and Aβ42 represent the major contributing factors; however, the inclusion of additional peptides further improves the model. Notably, the weightings were distinct when optimising for classification versus onset prediction, suggesting differential relevance of Aβ peptides. Finally, we observed that data from iPSC models show greater weighting for Aβ38 and less for Aβ40 compared with data from *PSEN1/2* double-knockout cell lines^8–10^.

Employing a patient-derived iPSC-neuronal model, this study complements previous studies in *PSEN1*/*PSEN2* double-knockout cell lines due to the presence of one healthy *PSEN1* allele and two healthy *PSEN2* alleles, analogous to the patient setting. This model therefore represents a physiological model of Aβ production in human neurons, which we have previously shown to faithfully correlate with clinical data from the same donor^6,13^. This model of Aβ production is relevant for current efforts in developing blood plasma biomarkers^23^.

Previously reported ratios performed well: Aβ37:42^8^ distinguished patient and control lines better than other linear ratios and Aβshort/long^9^ performed best for predicting age-at-onset. However, weighted combined value ratios (wCVR) were able to outperform linear ratios in both metrics, suggesting that increasing the number of Aβ peptides and assigning weight to each is able to further improve biomarker performance. The optimal ratio for all data, centred around both disease classification and predicting age-at-onset is: wCVR = (21 · Aβ37 + 10 · Aβ38 + 69 · Aβ40)/(94 · Aβ42 + 6 · Aβ43). Data-driven analyses support the increased pathogenicity of Aβ42 and Aβ43 versus Aβ37, Aβ38 and Aβ40; supporting in vitro data such as the differential aggregation kinetics of the longer species compared to the shorter species^24^.

A diversity of Aβ peptides exist, and EOFAD mutations alter their relative abundance differently. We analysed wCVR to infer the relative contribution of each peptide to classifying disease and for predicting age at onset. The fact that weights were dissimilar for the two biomarker analyses suggests that ‘pathogenicity’ and ‘severity’ show some distinction, for example changes to Aβ38 can inform on onset prediction but is negatively associated with classification. Additionally, we observed differences in the wCVR in iPSC-models compared to cell models, such that Aβ38 has a higher weighting at the expense of Aβ40. This may inform on the presence of a healthy *PSEN1* allele in iPSC models, speaking to the overrepresentation of Aβ40 compared with other peptides.

*PSEN1* mutation carriers show a diversity in the onset of fAD, even within families with the same mutation. Our analyses find no association between the location of the mutation across the PSEN1 protein and age-at-onset, similar to previous observations^10^. These data suggest that the severity of each mutation depends upon the biochemical effect of each amino acid substitution on the tertiary/quaternary protein structure.

Our study comes with some limitations to consider. One limitation is the low sample sizes. More donors and data replicates will further improve the confidence in these findings, for which replication studies are required. Another limitation is that the panel of iPSC samples tested comprise an over-representation of mutations that produce relatively high levels of Aβ43 (R278I^13^, E280G^14^ and P436S^15^), potentially skewing the relative contribution of Aβ43. However, our analysis of other datasets which do not have the same issue mitigates this limitation. Finally, age-at-onset is known to be variable and so the regression analyses should be considered with this caveat in mind.

In conclusion, this study presents new data, a new fluid biomarker harmonisation method, and a new ratio-based biomarker. Our experimental results demonstrate superior performance to the previous state-of-the-art ratio biomarkers, which have demonstrated that Aβ ratios correlate to measures of cognition^9,10^. Together, we have shown that our new approaches to harmonisation and for incorporating extra peptides in the ratio can contribute to the diagnosis of novel EOFAD mutations and potentially make plasma Aβ biomarker tests a clinical reality^23^.

## 6. Acknowledgements

The authors would like to acknowledge UCL Genomics for their support in the low depth whole genome sequencing analysis for karyotype stability assays.

---

**Supplementary Figure 1. Characterisation of newly generated *PSEN1* E184D iPSC line.** A) Immunocytochemistry of iPSCs for pluripotency markers NANOG and SSEA4. B) Sanger sequencing to confirm the point mutation Chr14:73659355 A>C. C) qPCR data of commonly duplicated or deleted genomic regions to support stable karyotype. D) Low coverage whole genome sequencing to confirm karyotype stability.

**Supplementary Figure 2. Characterisation of novel A**b**37 ELISA.** A) Linearity is shown using a serial dilution of recombinant Aβ37. The matrix (iPSC-neuron conditioned cell culture media) does not affect linearity of recombinant Aβ standards, evidenced by all datapoints raised by the same degree (corresponding to Aβ37 in the conditioned media). B) Representation of matrix effect plotted as a % recovery. C) Measures of specificity against other recombinant Aβ peptides. The Ab37 antibody shows some cross-reactivity with Aβ38, however this is 34-fold lower (right shift). D) Comparison of Ab37 quantification relative to Aβ42 from newly generated ELISA data with published mass spec data^13^.

**Supplementary Figure 3. Peptide abundance profiles.** Peptide abundance (fraction) per cohort (in grayscale) is shown on the vertical axis for Ab37, Ab38, Ab40, Ab42, and Ab43. UCL data had an over-representation of Aβ38 compared to the other cohorts.

**Supplementary Figure 4. Association of mutation with AAO and A**b**42:40 for all harmonised data.** Left: AAO (y-axis) plotted against the position of the mutation in PSEN1 (x-axis). Dashed line shows line of best fit. Right: Aβ42:40 (y-axis) plotted against position of the mutation in PSEN1 (x-axis). Dashed line depicts control Aβ42:40.

**Supplementary Figure 5. Peptide importance in wCVR.** The spread of peptide weights (between 0 and 100%) across 200 bootstrap experiments, signed to indicate numerator (+) or denominator (–) for three configurations of wCVR. When driven by R^2^ with AAO (Fitness = R2, upper right panel), wCVR ➔ Ab40/bB42 with some Ab37 in the denominator. When driven by precision-recall AUC for case-control classification (Fitness = PR_AUC, lower right panel), wCVR ≈ Aβshort/long. When driven by both metrics, wCVR is a hybrid: dominated by Aβ40/AB42, but with Aβ37 in the numerator.

**Supplementary Figure 6. Biomarker performance by peptide weight (bootstrap).** Biomarker performance (vertical axis) as a function of peptide weight (horizontal axis). Each dot represents one of 200 bootstrap experiments.

**Supplementary Figure 7. Biomarker performance by peptide weight (grid search).** Left: wCVR R^2^ with AAO. Right: precision-recall AUC. The colour gradient shows a sweet spot where biomarker performance is improved by adding other peptides to either Ab40/Ab42 (left) or Ab37/Ab42 (right).

