## Supplementary figures for "Unbiased data-driven analysis of five amyloid-beta peptides for biomarker investigations in familial Alzheimer’s disease"

### Slide 1
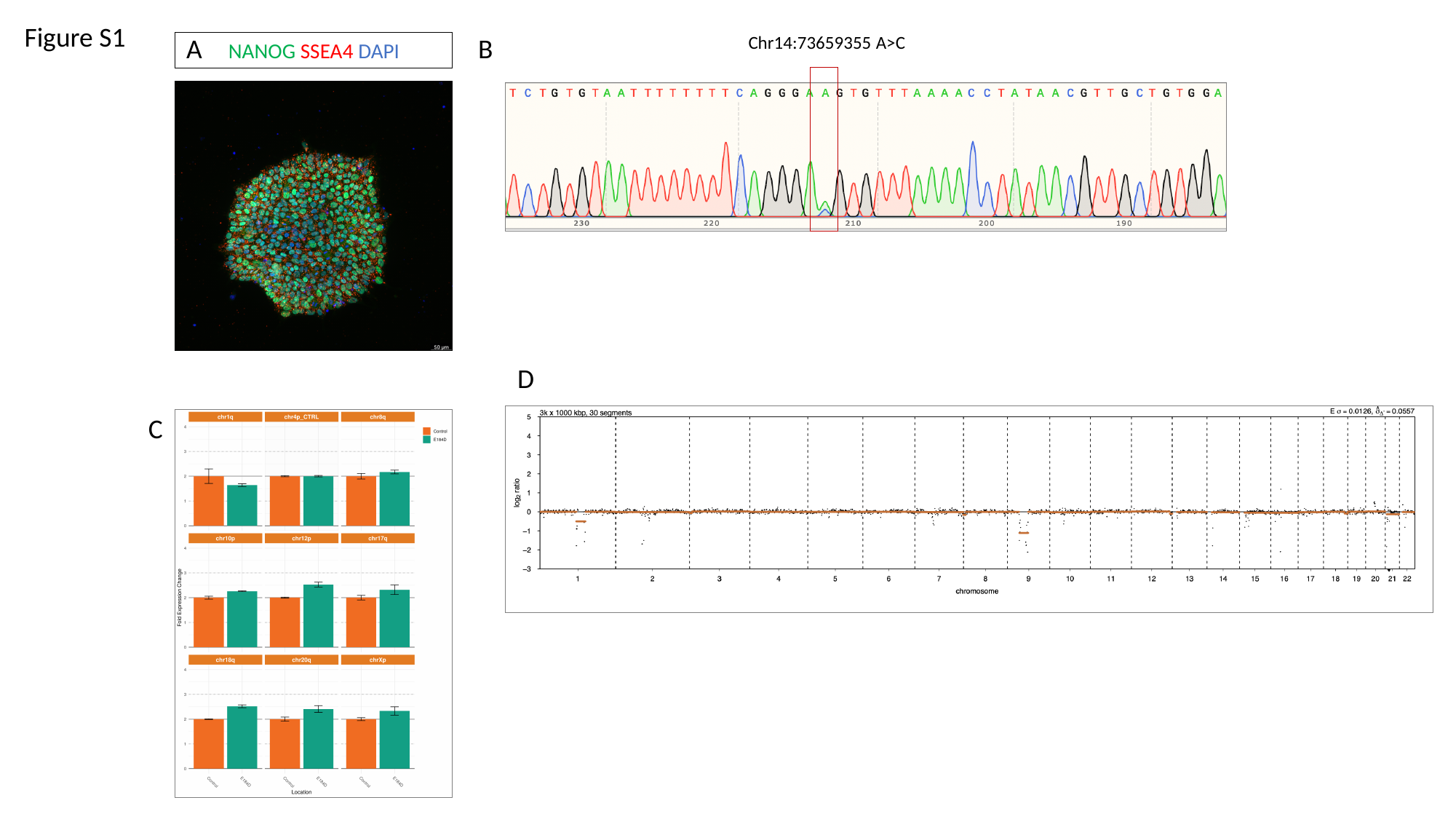

Figure S1
A
B
 Chr14:73659355 A>C
NANOG SSEA4 DAPI
D
C

### Slide 2
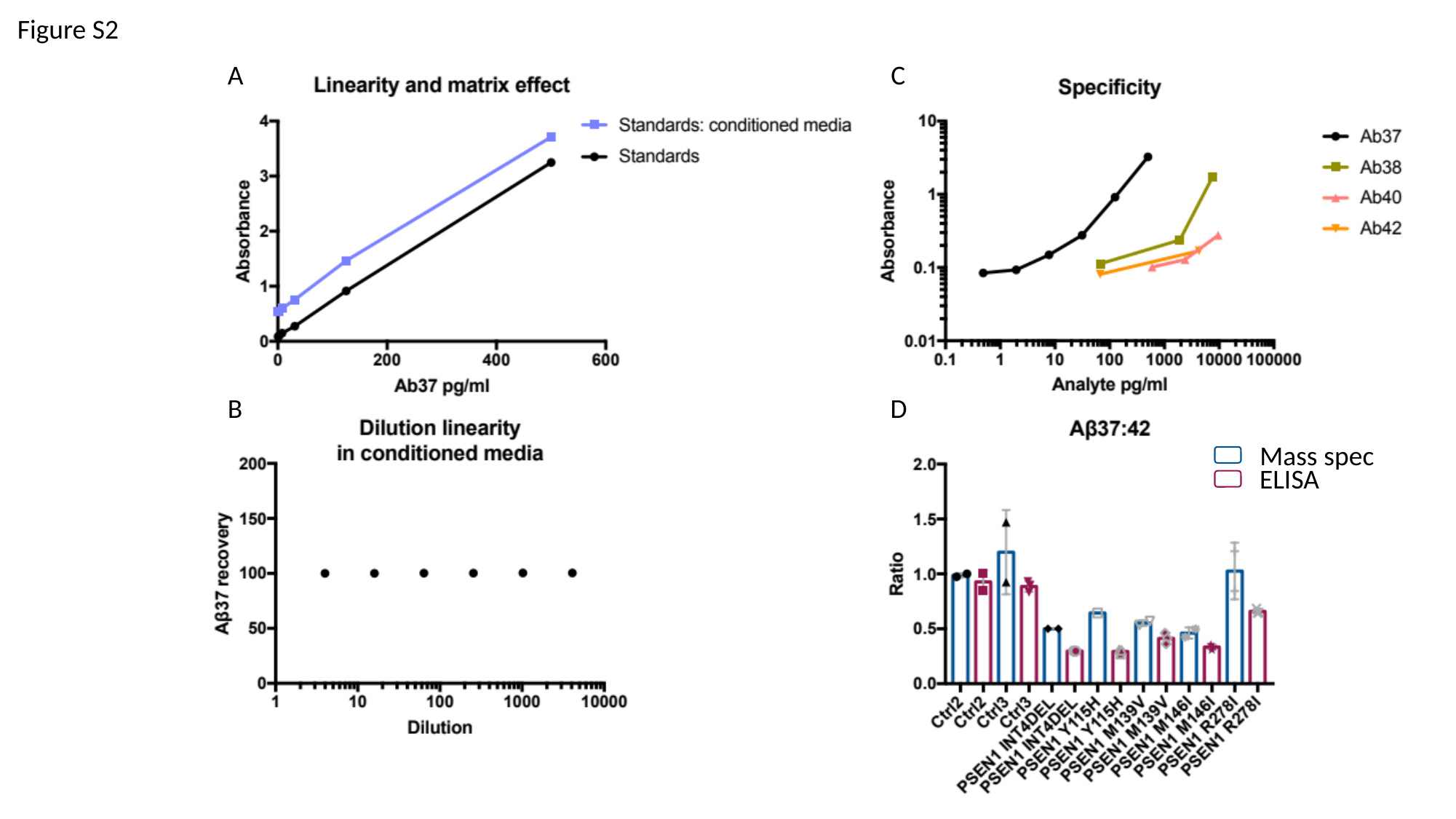

Figure S2
A
C
B
D
Mass spec
ELISA

### Slide 3
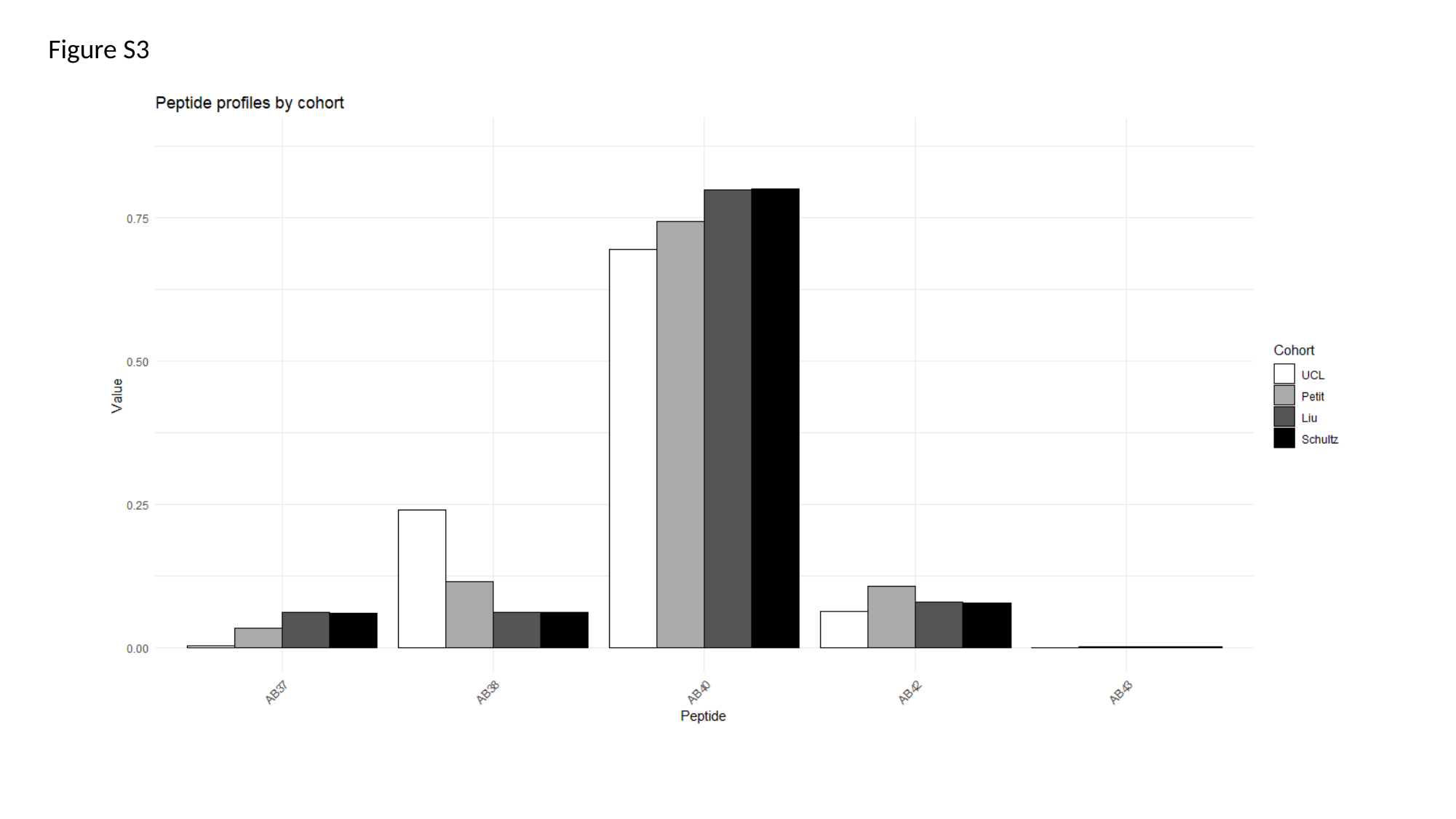

Figure S3

### Slide 4
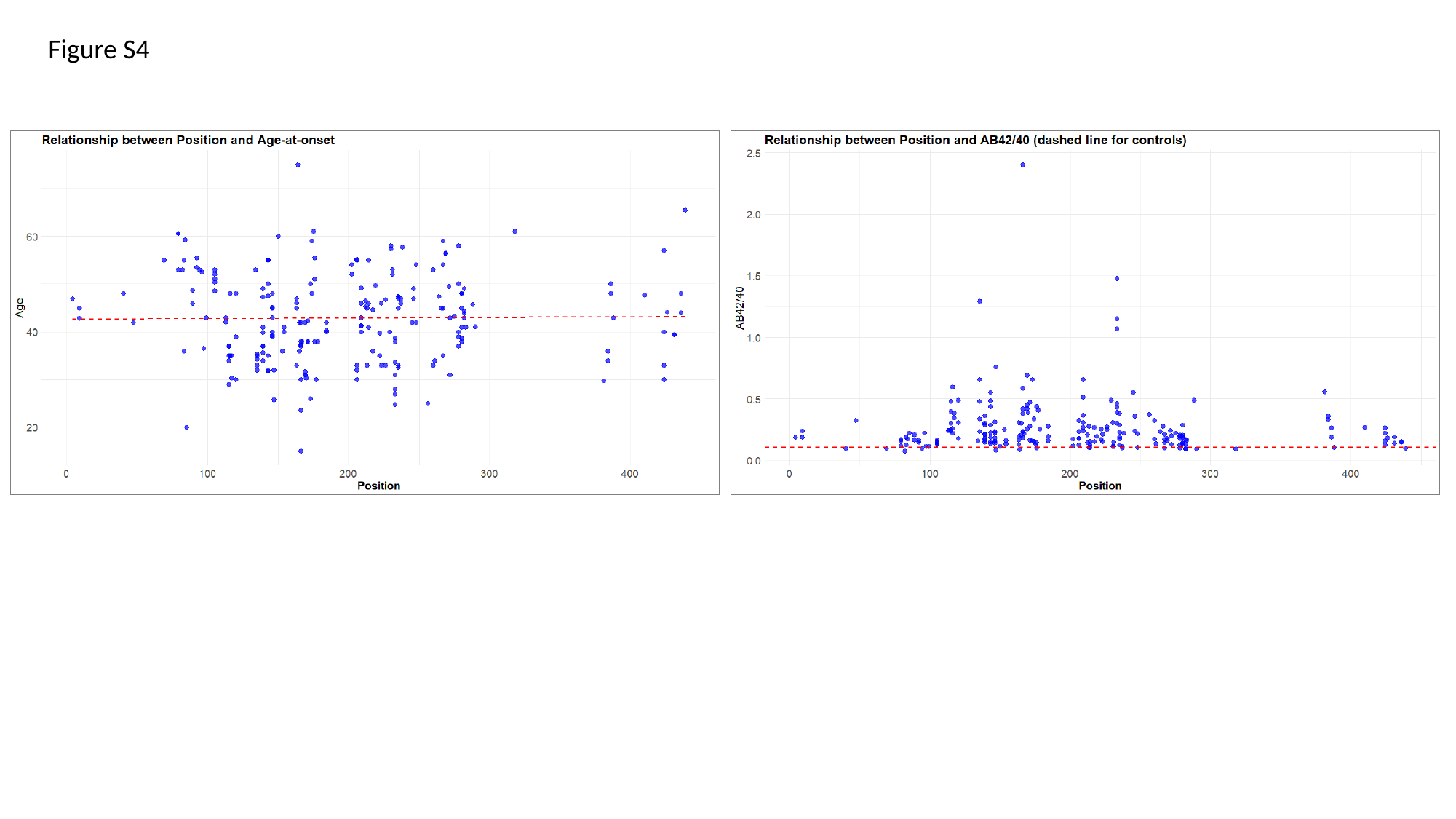

Figure S4

### Slide 5
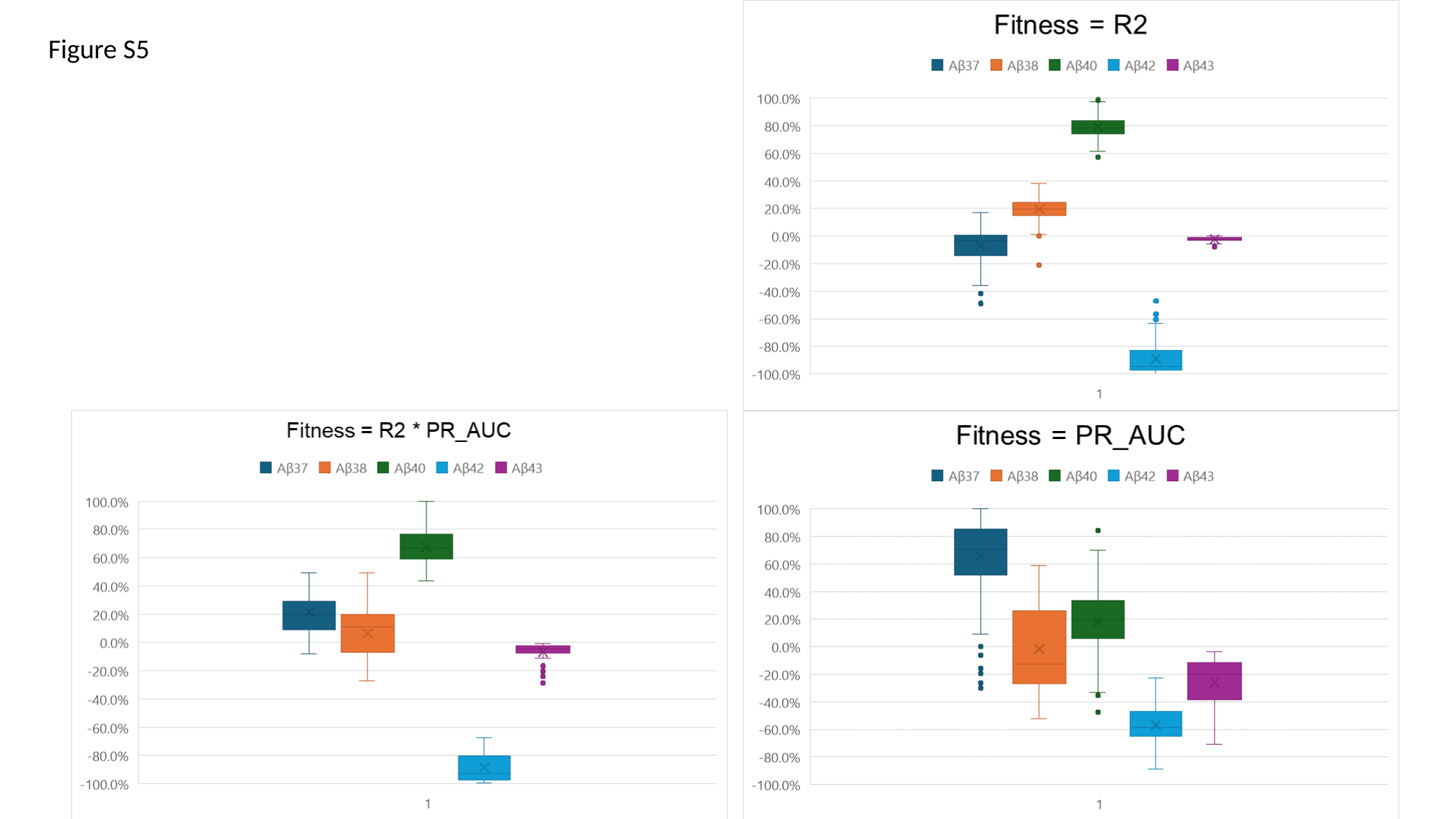

Figure S5

### Slide 6
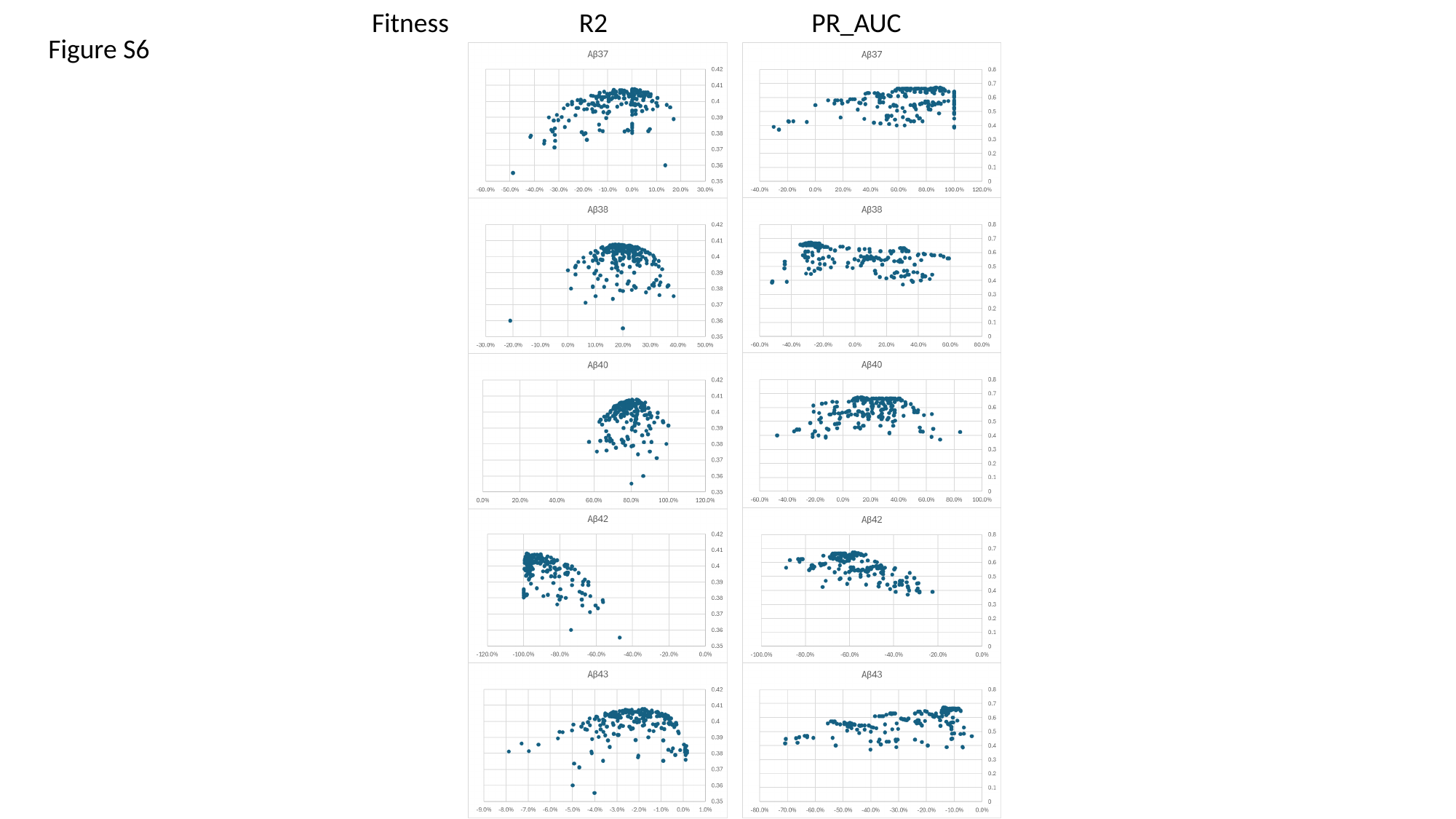

Fitness R2 PR_AUC
Figure S6

### Slide 7
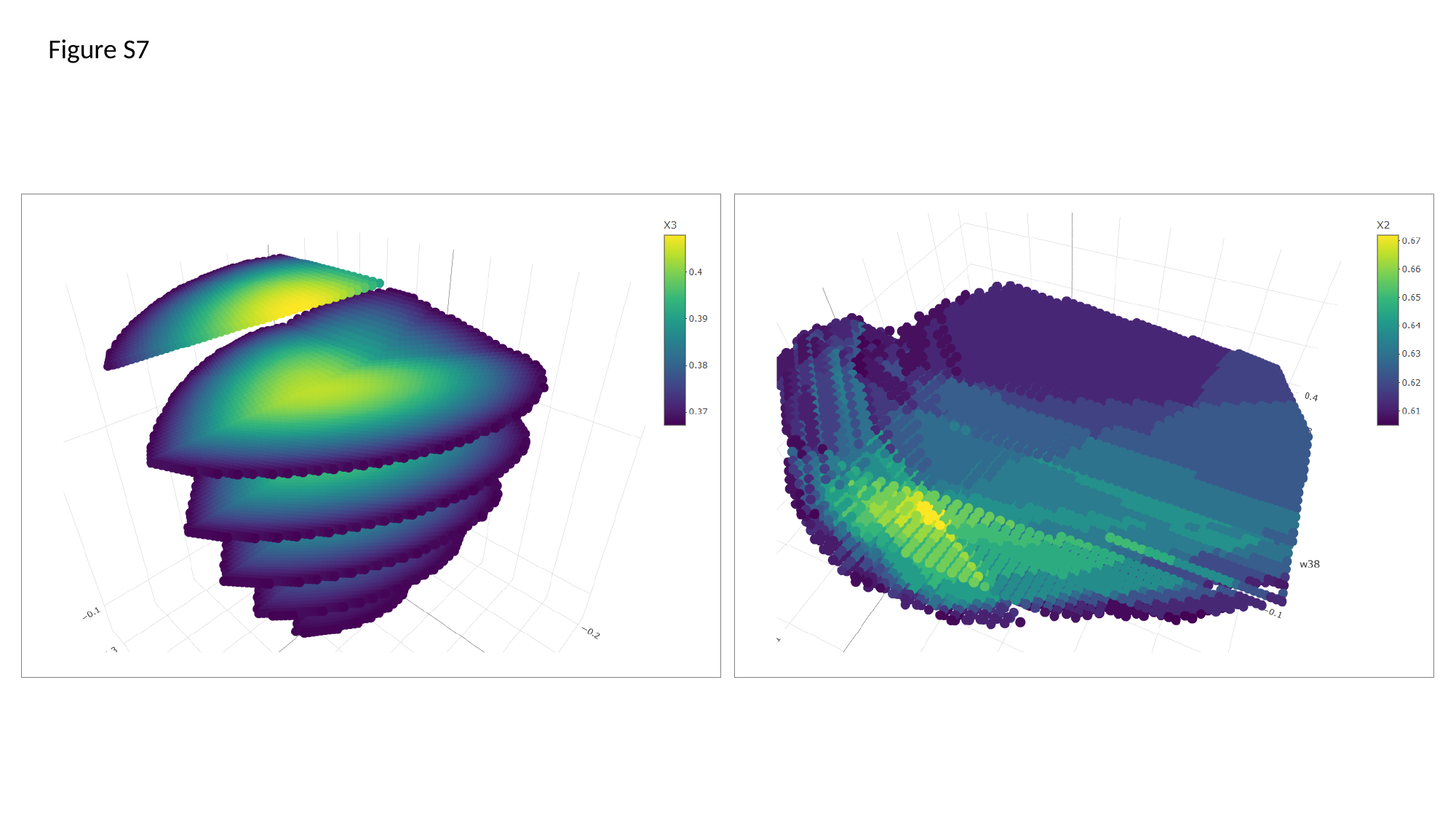

Figure S7
