## Supplementary material for "Unbiased data-driven analysis of five amyloid-beta peptides for biomarker investigations in familial Alzheimer’s disease": Tables 2-4 and Table S1

**Table 2: Correlation measures, per peptide, per cohort (and all data “All”), with age-at-onset**. In bold, uncorrected p-values below 5%.

|  |  | **Pearson** | | | |  | **Kendall** | |
| --- | --- | --- | --- | --- | --- | --- | --- | --- |
|  | **Peptide** | **ρ** | **p-value** | **Lower_CI** | **Upper_CI** |  | **τ** | **p-value** |
| **iPSC** | Aβ37 | 0.114 | 0.754 | -0.555 | 0.694 |  | -0.068 | 0.787 |
|  | Aβ38 | -0.156 | 0.666 | -0.716 | 0.525 |  | -0.068 | 0.787 |
|  | Aβ40 | 0.462 | 0.179 | -0.237 | 0.846 |  | 0.296 | 0.241 |
|  | Aβ42 | -0.559 | 0.093 | -0.879 | 0.109 |  | -0.341 | 0.176 |
|  | Aβ43 | 0.358 | 0.310 | -0.351 | 0.806 |  | 0.159 | 0.528 |
| **Petit** | Aβ37 | -0.141 | 0.512 | -0.515 | 0.279 |  | -0.167 | 0.268 |
|  | Aβ38 | -0.093 | 0.666 | -0.478 | 0.323 |  | -0.022 | 0.902 |
|  | Aβ40 | **0.781** | **0.000** | 0.551 | 0.900 |  | 0.534 | **0.000** |
|  | Aβ42 | **-0.834** | **0.000** | -0.926 | -0.650 |  | -0.710 | **0.000** |
|  | Aβ43 | -0.292 | 0.167 | -0.622 | 0.127 |  | -0.241 | 0.101 |
| **Liu** | Aβ37 | **0.253** | **0.004** | 0.085 | 0.407 |  | 0.172 | **0.004** |
|  | Aβ38 | **0.194** | **0.026** | 0.024 | 0.354 |  | 0.190 | **0.001** |
|  | Aβ40 | **0.486** | **0.000** | 0.343 | 0.607 |  | 0.342 | **0.000** |
|  | Aβ42 | **-0.641** | **0.000** | -0.732 | -0.528 |  | -0.521 | **0.000** |
|  | Aβ43 | **-0.238** | **0.006** | -0.394 | -0.070 |  | -0.144 | **0.016** |
| **Schultz** | Aβ37 | 0.043 | 0.751 | -0.222 | 0.303 |  | 0.092 | 0.319 |
|  | Aβ38 | -0.007 | 0.958 | -0.270 | 0.256 |  | 0.187 | **0.043** |
|  | Aβ40 | **0.467** | **0.000** | 0.233 | 0.650 |  | 0.334 | **0.000** |
|  | Aβ42 | **-0.581** | **0.000** | -0.732 | -0.376 |  | -0.430 | **0.000** |
|  | Aβ43 | 0.001 | 0.995 | -0.262 | 0.264 |  | 0.063 | 0.497 |
| **All** | Aβ37 | 0.031 | 0.644 | -0.101 | 0.163 |  | 0.074 | 0.104 |
|  | Aβ38 | 0.066 | 0.330 | -0.067 | 0.196 |  | 0.103 | **0.024** |
|  | Aβ40 | **0.493** | **0.000** | 0.386 | 0.587 |  | 0.343 | **0.000** |
|  | Aβ42 | **-0.580** | **0.000** | -0.661 | -0.485 |  | -0.447 | **0.000** |
|  | Aβ43 | -0.108 | 0.109 | -0.237 | 0.024 |  | -0.095 | **0.038** |

**Table 3: Evaluation metrics for the biomarkers considered.** In bold, the best result per metric. wCVR = weighted composite value ratio, which is a linear combination of peptides both in the numerator and denominator. The suffix -R indicates that the wCVR was optimised for R^2^ with respect to age-at-onset; -P indicates that wCVR was optimised to maximize the precision-recall area-under-the-curve for the controls; -PR indicates that wCVR was optimised to maximize the product of both metrics (regression with age-at-onset and classification for the controls). AUC = area under the receiver operator characteristic curve. PR AUC = area under the precision-recall curve. fAD = data from familial Alzheimer’s disease (mutation-carriers). R^2^ = coefficient of determination, using Pearson’s correlation with age-at-onset.

| **Biomarker** | **AUC** | **PR AUC controls** | **PR AUC fAD** | **R^2^** |
| --- | --- | --- | --- | --- |
| wCVR-R | 0.977  (0.953, 0.994) | 0.456  (0.287, 0.805) | 0.999  (0.998, 1.000) | **0.412  (0.284, 0.526)** |
| wCVR-RP | 0.986  (0.966, 0.997) | 0.593 (0.363, 0.934) | 0.999  (0.999, 1.000) | 0.380  (0.255, 0.497) |
| wCVR-P | **0.992  (0.977, 1.000)** | **0.700  (0.435, 1.000)** | **1.000 (0.999, 1.000)** | 0.242  (0.145, 0.350) |
| Aβ42/40 | 0.961  (0.929, 0.983) | 0.345  (0.217, 0.615) | 0.998  (0.997, 0.999) | 0.259  (0.187, 0.343) |
| Aβ37/42 | 0.977  (0.953, 0.993) | 0.454  (0.29, 0.785) | 0.999  (0.998, 1.000) | 0.114  (0.048, 0.230) |
| short/long | 0.974  (0.947, 0.991) | 0.411  (0.265, 0.729) | 0.999  (0.998, 1.000) | 0.403  (0.275, 0.517) |

**Table 4: wCVR weights per configuration (all data).**

Suffix -R optimizes wCVR for R^2^ versus age-at-onset. Suffix -P optimizes wCVR versus precision-recall area under the curve (PR AUC). Suffix -RP optimizes for the multiplication of both R^2^ and PRAUC. wCVR = weighted composite value ratio, which is a linear combination of peptides both in the numerator and denominator.

| **A) Data-driven weighting and combination of peptides (wCVR)** | | | | | | | |
| --- | --- | --- | --- | --- | --- | --- | --- |
| **Aβ37** | **Aβ38** | **Aβ40** | **Aβ42** | **Aβ43** | **biomarker** | **PR AUC** | **R^2^** |
| **-0.2%** | 19.0% | 81.0% | **-97.8%** | **-2.0%** | wCVR-R | 0.456 | 0.412 |
| 21.0% | 9.8% | 69.2% | **-93.6%** | **-6.4%** | wCVR-RP | 0.593 | 0.382 |
| 88.9% | **-27.8%** | 11.1% | **-59.0%** | **-13.2%** | wCVR-P | 0.705 | 0.185 |
| **B) Short/long relative weights per cohort (pre-harmonisation)** | | | | | | | |
| **Aβ37** | **Aβ38** | **Aβ40** | **Aβ42** | **Aβ43** | **Cohort** |  |  |
| 0.3% | 25.7% | 74.1% | **-99.7%** | **-0.3%** | **iPSC** |  |  |
| 3.7% | 13.0% | 83.3% | **-99.4%** | **-0.6%** | **Petit** |  |  |
| 6.6% | 6.7% | 86.7% | **-98.6%** | **-1.4%** | **Liu** |  |  |
| 6.6% | 6.6% | 86.8% | **-98.6%** | **-1.4%** | **Schultz** |  |  |

**Table S1: Permutation feature importance analysis.** In bold, the peptide with the biggest effect. Both biomarkers (short/long and wCVR-RP) have a similar composition: the peptides Aβ37, 38 and 40 in the numerator, and Aβ42 and 43 in the denominator.

|  | **Feature importance (90% confidence interval)** | | | |
| --- | --- | --- | --- | --- |
|  | **1 - Precision-recall AUC** | | **1 - (R2 vs AAO)** | |
|  | **short/long** | **wCVR-RP** | **short/long** | **wCVR-RP** |
| Aβ37 | 0.99 (0.98, 1.04) | 1.13 (1.02, 1.19) | 1.00 (0.99, 1.00) | 0.99 (0.97, 1.02) |
| Aβ38 | 1.10 (1.01, 1.17) | 1.09 (1.06, 1.19) | 1.05 (1.02, 1.09) | 1.03 (1.00, 1.05) |
| Aβ40 | 1.06 (0.92, 1.14) | 1.18 (1.00, 1.24) | 1.06 (1.05, 1.09) | 1.06 (1.05, 1.08) |
| **Aβ42** | **1.58 (1.56, 1.61)** | **1.99 (1.82, 2.09)** | **1.61 (1.54, 1.66)** | **1.52 (1.45, 1.57)** |
| Aβ43 | 1.02 (1.01, 1.19) | 1.55 (1.18, 1.57) | 1.03 (1.00, 1.05) | 1.19 (1.10, 1.22) |
